# Microsporidia dressing up: the spore polaroplast transport through the polar tube and transformation into the sporoplasm membrane

**DOI:** 10.1101/2023.05.01.538940

**Authors:** Qing Lv, Yuqing Chen, Lei Qi, Liuyi Hong, Zhengkai Xie, Hongjie Liao, Xiuli He, Chenxi Li, Xianzhi Meng, Jie Chen, Bing Han, Qingtao Shen, Louis M. Weiss, Zeyang Zhou, Mengxian Long, Guoqing Pan

## Abstract

Microsporidia are obligate intracellular parasites that infect a wide variety of hosts, including humans. Microsporidian spores possess a unique, highly specialized invasion apparatus involving the polar filament, polaroplast and posterior vacuole. During spore germination, the polar filament is discharged out of the spore forming the hollow polar tube that transports the sporoplasm components including nucleus into the host cell to achieve the invasion. Due to the complicated topological changes occurring in this process, the formation of sporoplasm is unclear. Here, electron microscopy observation and DiI staining confirmed that during spore germination, a large number of vesicles derived from the polaroplast, nucleus and other cytoplasm were transported out via the polar tube. Meanwhile, the posterior vacuole and plasma membrane remained in the empty spore coat. In addition, there was no DiI-labeled membrane around the nucleus in mature spores, whereas a DiI-labeled limit membrane wrapping nucleus was found at the tip of the extruded polar tube, suggesting that the membrane of sporoplasm was formed outside the mature spore. Two *Nosema bombycis* sporoplasm surface proteins (NbTMP1 and NoboABCG1.1) were located at the polaroplast in mature spores, in the extruded polar tube and on the sporoplasm membrane, which indicated that the polaroplast transported via the polar tube finally became the limiting membrane of the sporoplasm. Golgi-tracker green and Golgi marker protein syntaxin 6 were also found the same model, which was consistent with the transported polaroplast derived from Golgi transformed into the novel sporoplasm membrane during spore germination.

**Importance:** Microsporidia, obligate intracellular pathogenic organisms, cause huge economic losses in agriculture and even threaten human health. The key to successful infection of microsporidia is its unique invasion apparatus which includes the polar filament, polaroplast and posterior vacuole. When the spore is activated to geminate, the polar filament uncoils and undergoes a rapid transition into the hollow polar tube that will transport the sporoplasm components including nucleus into a host cell to achieve the invasion. Knowledge of structure difference between polar filament and polar tube, the process of cargo transport in extruded polar tube, and the formation of the sporoplasm membrane are still poorly understood. Herein, we verify that the polar filament evaginates to form the polar tube, which serves as a conduit for transporting elongated nucleus and other sporoplasm components. And we confirm that the transported polaroplast finally transforms into the novel sporoplasm membrane during spore germination. Our study provides new insights into the cargo transportation process of polar tube and origin of the sporoplasm membrane, which serve as foundations for clarifying the microsporidian infection mechanism.

## Introduction

Microsporidia are a large group of important parasites in agriculture and human health [1, 2]. *Nosema bombycis* was identified in 1857 as the cause of Pébrine in the silkworm *Bombyx mori* in Europe [3], since then over 200 genera and 1700 species of microsporidia have been identified [4–6]. These enigmatic pathogens have been fascinated for more than 160 years because of their wide range of hosts, from invertebrates to vertebrates, as well as their unique and highly specialized invasion apparatus [7–11]. Microsporidia not only cause huge economic losses to agricultural production [12], but also are responsible for human diseases. It was initially thought to be restricted to infect immune compromised humans; however, it is now known that the microsporidian infection can also occur in immune competent individuals [13, 14].

Microsporidia form characteristic infectious spores, which can survive outside their host cells [1, 11]. Mature spores have a unique infection apparatus for invasion, which includes the polar filament (defined as the polar tube after spore germination), polaroplast and posterior vacuole (PV) [11, 15, 16]. In all microsporidian mature spores [11, 17], the polar filament tightly coils and forms a spring-like structure [18], which is a solid structure comprising electron-dense alternating concentric rings in cross-sections [6, 19]. In 2022, the macrostructure of the polar filament in *Encephalitozoon hellem* was characterized by Cryo-EM and demonstrated that the polar filament had a multi-layer concentric circle structure and many bumps with a diameter of 2.5 nm were present on the second layer of polar filament [20]. The polaroplast is a system of membrane-limited cavities in the anterior part of the spore, which occupies one-third to one-half of the spore volume [11, 17]. The compartments of the polaroplast are shaped like lamellae, chambers, or tubules delimited by approximately 5 nm thick unit membranes [11, 21]. The posterior polaroplast is closely to the linear portion of the polar filament to stabilize the polar filament. The posterior vacuole (PV) at the end of the spore is roughly bowl-shaped [6, 21, 22], which may also play a vital role in spore germination, providing dynamic support for polar tube extension and cargo rapid passage [1, 22–24]. The PV membrane is interlaced with the polar filament coils, suggesting that it may interact with the polar filament [21].

Under an appropriate stimulus spore germination occurs and the polar filament everts from the thinnest point of the spore forming a hollow tube, the polar tube [6, 11]. At present, six polar tube proteins (PTPs) have been identified in a variety of microsporidia [12, 25–30], and many of them appear to be involved in polar tube attachment to host cell and interaction with host [6, 11, 12]. Till now, the evagination mechanism of the polar filament as well as the detailed infection process of microsporidia has not been clarified [14, 31–33]. It has been suggested that the polar filament everts during germination like “reversing finger of a glove”. During spore germination, the enlargement of polaroplast and the expansion of PV increase the internal turgor pressure of the spore to promote polar tube release and push the spore contents into the polar tube [3, 6, 11].

The sporoplasm is indispensable for microsporidian infection and proliferation. Once the polar tube is released, the infectious cargo, including the nucleus, is transported through the polar tube to form sporoplasm. The sporoplasm then appears as a droplet at the distal end of the polar tube and remains attached with the polar tube for several minutes [7, 13]. Interaction between the polar tube and sporoplasm remain to be defined [1, 6, 34, 35]. Studies have shown that PTP4 from *Encephalitozoon hellem* interacts with the sporoplasm surface protein 1 (SSP1), for assisting the sporoplasm attached with the tip of polar tube [6]. The sporoplasm released from polar tube has a limiting membrane derived from polaroplast [11, 36–38]. However, there is no conclusive evidence to confirm this hypothesis. In this study, we verified that the extruded polar tube served as a conduit for transporting elongated nucleus and sporoplasm components. And we confirmed that the transported polaroplast finally transformed into the novel sporoplasm membrane after spore germination.

## Results

### Process of polar tube germination and cargo transportation

Although the infection mechanism of microsporidia is still unclear, it is indisputable that spore germination and polar tube ejection are essential [1, 6, 11]. We found that the extruded polar tube of *N. bombycis* was about 150 μm in length, nearly 50 times the length of the spore by TEM. Discontinuous high electron density material was observed in the polar tube that are likely intracellular cargo traversing the polar tube (Fig. S1A). In addition, curved hooks were frequently observed at the anterior end of extruded polar tube (Fig. S1A and S1B). Further, the *N. bombycis* spores were stimulated to germinate on the Cryo-EM grid, which were then rapidly frozen by liquid nitrogen and ethane. Cryo-EM results demonstrated a large number of germinated spores and extruded polar tubes on the grid (Fig. 1A). Two types of polar tube were observed. One type was uniform with about 2 nm long bumps distributed on its surface (Fig. 1B). The other type demonstrated a variety of lumens and vesicles distributed discontinuously within the polar tube. These cargoes had different shapes, such as elliptic, long strips and rounded and most of them were monolayer membrane structures. The surface of these polar tubes was no longer regular and was covered with relatively loose fibrillar material, which may extend as far as 2-15 nm from the polar tube surface (Fig. 1C and 1D). The diameter of polar tubes without vesicle (type Ⅰ) was 80-110 nm, whereas the diameter of polar tubes containing vesicles (type Ⅱ) varied greatly, ranging from 120-205 nm (Fig. 1E), indicating that the polar tube was an elastic tubular structure with the expansion capacity for cargo passage.

**Fig. 1.**
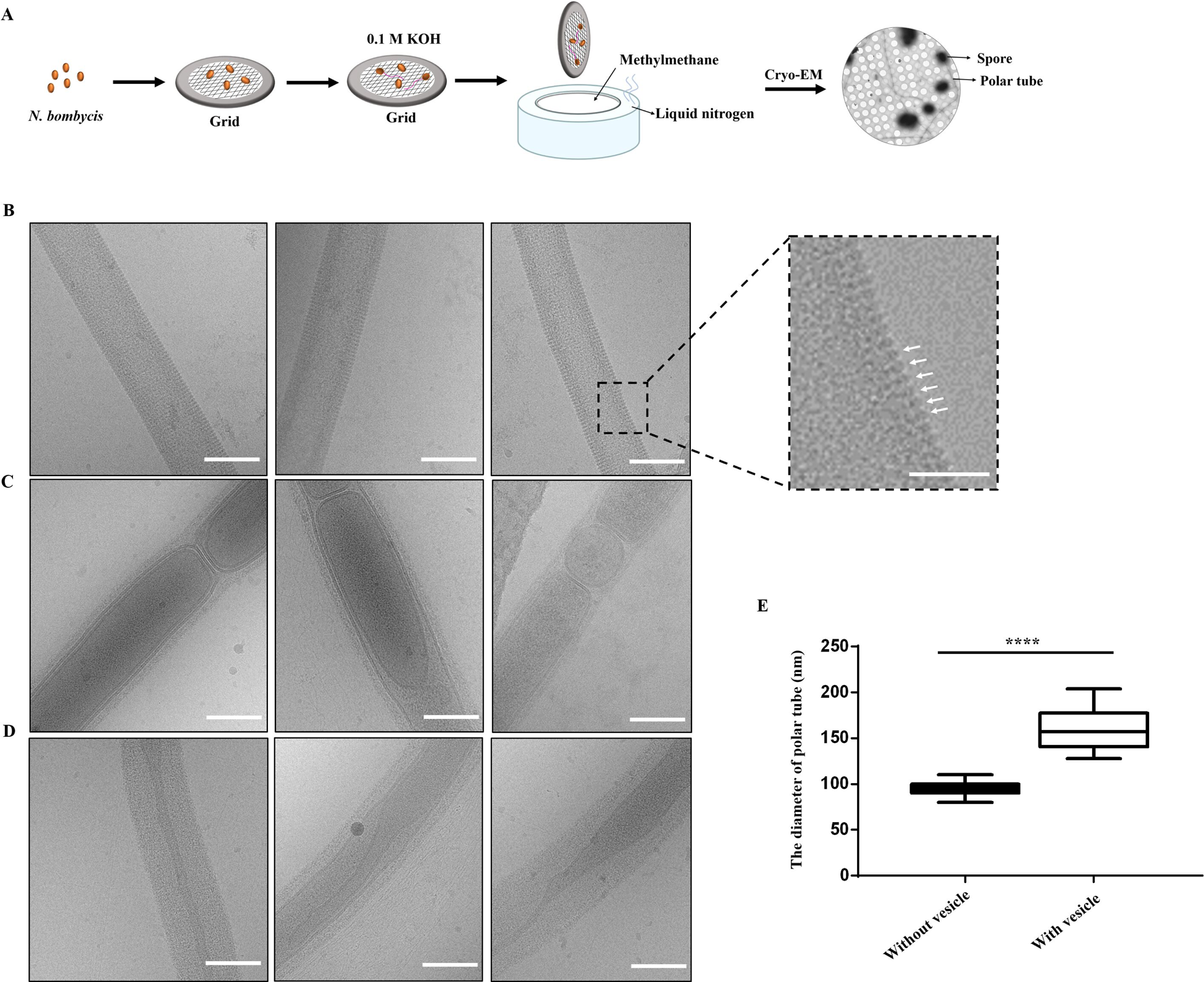
Cryo-EM analysis of the extruded polar tube of *N. bombycis*. (A) Schematic of the general workflow for Cryo-EM. (B-D) Cryo-EM analysis of the different structural characteristics of the polar tube. *Scale-bars*: 100 nm. The white arrow in the enlarged image pointed to the bumps distributed on surface of polar tube. *Scale-bars*: 50 nm. (E) Comparison of the diameter of polar tube with and without vesicle. (100 polar tube samples with and without vesicle were respectively selected and measured; ****, *p*<0.0001).

A lipophilic fluorescent dye DiI and the nucleus dye DAPI were used to label the cargo in the polar tube. We found that two nuclei of *N. bombycis* were elongated and deformed when they entered into the polar tube (Fig. 2A and 2B), and then returned to a spherical shape after reaching the tip of the polar tube (Fig. 2C). In addition, the red fluorescence signal of DiI appeared discontinuously throughout polar tube before, during and after the nuclei transportation, implying that besides the nuclei, polar tube also transported some DiI-labeled discontinuous membrane structure, which was consistent with Cryo-EM observation. More importantly, as shown in Fig. 2A and 2B, the nuclei in the polar tube was not surrounded by the DiI-labeled membrane. However, when the nucleus reached the tip of the polar tube, they were wrapped by the DiI-labeled membrane and adhered with the tip of the polar tube (Fig. 2C).

**Fig. 2.**
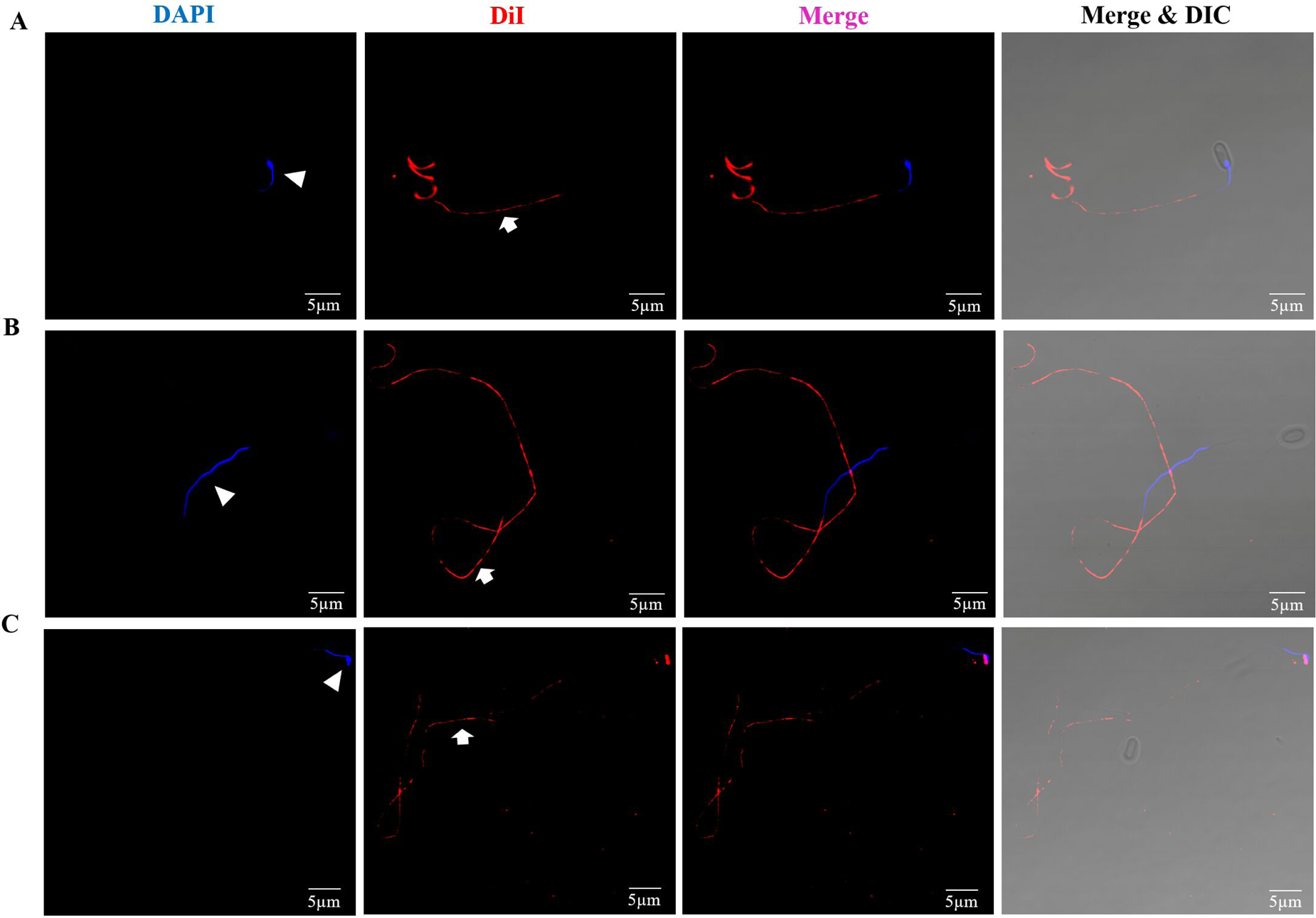
Fluorescent analysis of the cargo transport process through the polar tube of *N. bombycis*. (A) The polar tube was stained with DiI (red) before the nucleus (blue) was transported. (B) The polar tube was stained with DiI (red) during the nucleus (blue) was transported. (C) The polar tube was stained with DiI (red) after the nucleus (blue) was transported. The white triangle and arrow represented the nucleus and polar tube respectively. *Scale-bars*: 5 μm.

### The spore polaroplast transport via polar tube during spore germination

The fine structure of the polar filament in the mature spore was analyzed to determine whether these membrane structures were present in the polar filament prior to the germination. However, the purified polar filament fragment of *N. bombycis* was not labeled by DiI (Fig. S2A), which confirmed that the vesicle appeared in polar tube were absent in polar filament. By Cryo-EM, we found there were continuous and relatively high electron density material in the center region of polar filament, meanwhile, no vesicle was observed inside the polar filament (Fig. S2B). Therefore, we speculated that these DiI-labeled membrane structures existed in mature spores and entered into the polar tube for transportation during spore germination. Hence, the DiI was used to label the mature spore, and red fluorescence signal was identified at the location of polaroplast and PV in the mature spore. Meanwhile, the plasma membrane of the spores was also partially stained (Fig. 3A). A red ring was observed in the empty spore coats after spore germination (Fig. 3B), which was markedly different from that observed in mature spores. It was suggested that part of membrane structure in the mature spore was transported out during spore germination while the rest remained in the empty spore coat.

**Fig. 3.**
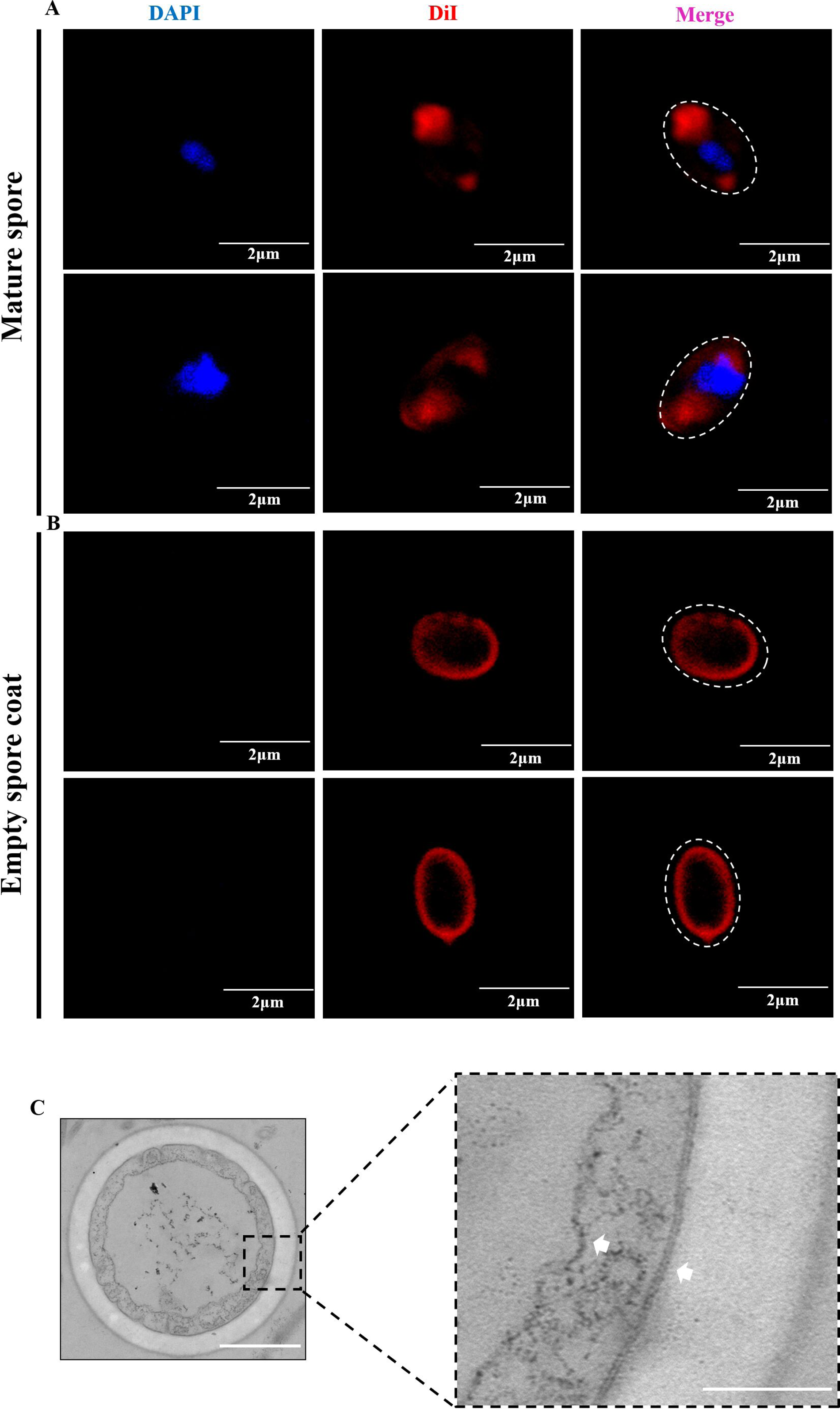
Analysis the membrane structure in the mature spore and empty spore coat of *N. bombycis*. (A) Fluorescence analysis the membrane structure in the mature spore by DiI staining. (B) Fluorescence analysis the membrane structure in the empty spore coat after germination by DiI staining. The nuclei were stained with DAPI (blue), and the membrane was stained with DiI (red). *Scale-bars*: 2 μm. (C) TEM observation the cross-section of the empty spore coat. *Scale-bars*: 500 nm. The white arrows were shown the PV membrane and the plasma membrane in the larger version of the black rectangle. *Scale-bars*: 200 nm.

Next, germinated and ungerminated of *N. bombycis* spores were sectioned and observed by TEM. In the ungerminated mature spores, the polar filament, the polaroplast and the PV were arranged in an orderly status (Fig. S3A). During spore germination, the polar filament and polaroplast were discharged through the polar tube, and then the PV gradually expanded to the volume of the spore and remained in the empty spore coat (Fig. S3B-S3E), which was consistent with previous results [3, 11]. Transverse section of empty spore coat demonstrated two membranes remained in empty spore coat after germination, the plasma membrane and PV membrane respectively. It is worth noting that the plasm membrane was a typical phospholipid bilayer structure, while the PV membrane was obviously different, and the structure was loose and irregular (Fig. 3C). Therefore, the membrane structure in the extruded polar tube were most likely the polaroplast. On the other hand, the area circled by the white dotted line clearly showed the characteristics of the polaroplast in mature spores, with a large number of sac or lamellae piled together (Fig. S3F), which were consistent with the vesicle membrane structures in the polar tube (Fig. 1C and 1D). Based on above results, we verified that these DiI-labeled vesicles in the polar tube were the transported polaroplast during spore germination.

### The process of the sporoplasm formation

The sporoplasm formation process after *N. bombycis* spore germination was observed *in vitro*. After 0.1 M KOH treatment for 5 minutes, the transported nuclei were wrapped in DiI-labeled membrane at the anterior end of extruded polar tube. At this time, the sporoplasm attached with the polar tube was irregular in shape, and their sizes were not consistent (Fig. 4A). About 7 minutes for germination later, most sporoplasms were detached from the tip of polar tube. And the shape of sporoplasm was still irregular, but tended to be round with almost 2 μm in diameter (Fig. 4B). About 10 minutes for germination later, the sporoplasm became spherical, exceeded 2 μm in diameter, and the nuclei were located near the inner edge of the sporoplasm membrane (Fig. 4C). Besides that, we also observed spontaneous spore germination process in *N. bombycis*-infected BmE cells and *E. hellem*-infected Vero E6 cells. The result was consistent with the observation of spore germination *in vitro*, which was the transported nuclei surrounded by the DiI-labeled membrane derived from polaroplast formed the sporoplasm at the tip of the polar tube (Fig. 4D and 4E).

**Fig. 4.**
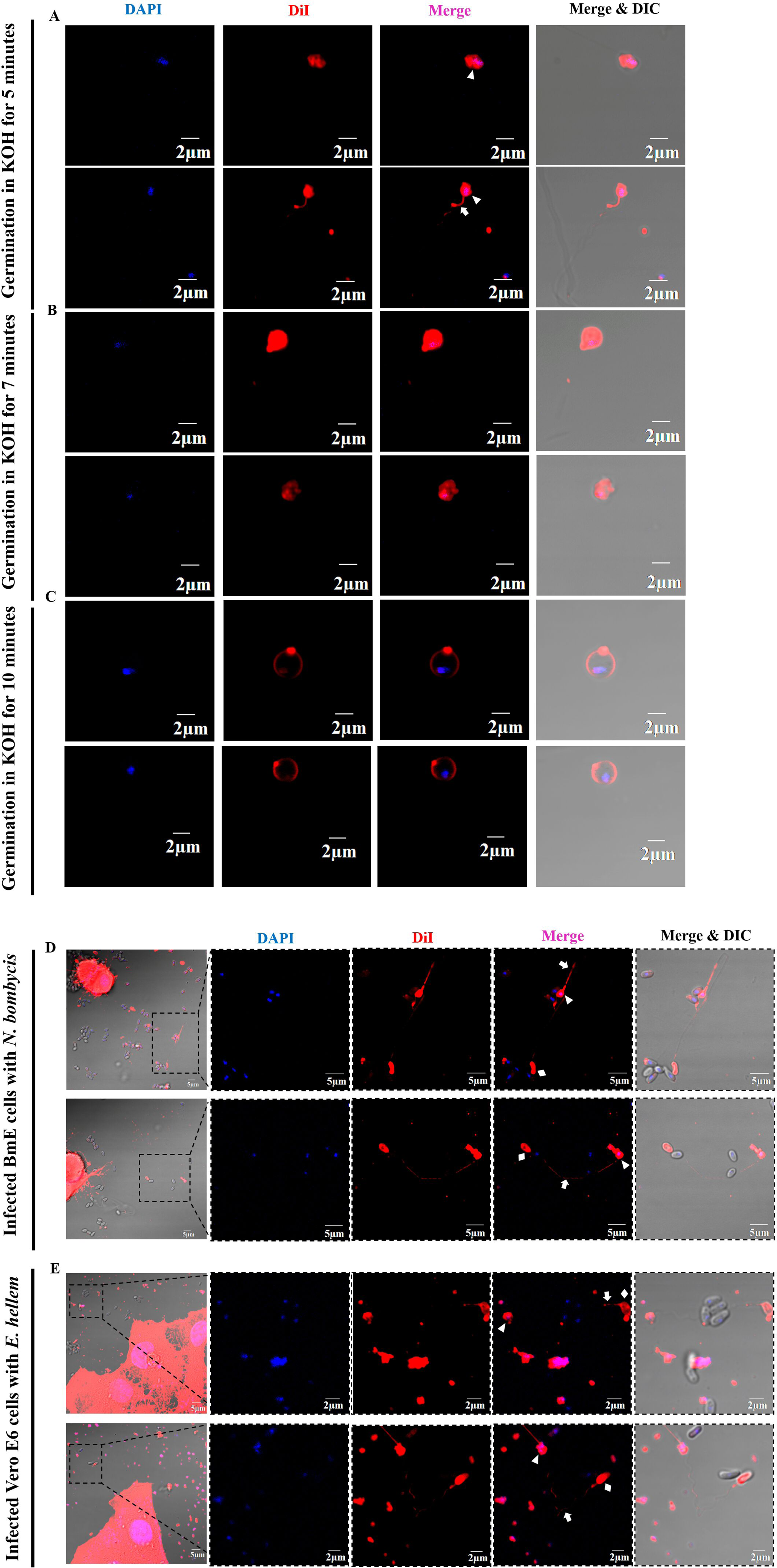
Analysis of the sporoplasm formation process. (A) Fluorescence analysis of the spore germination in KOH for 5 min. The sporoplasm was located at the tip of the polar tube. (B) Fluorescence analysis of the spore germination in KOH for 7 min. The sporoplasm was dropped from the polar tube with irregular shape. (C) Fluorescence analysis of the spore germination in KOH for 10 min. The sporoplasm was free with spherical shape. *Scale-bars*: 2 μm. (D) Fluorescence analysis of *N. bombycis*-infected BmE cells. *Scale-bars*: 5 μm. (E) Fluorescence analysis of *E. hellem*-infected Vero E6 cells. *Scale-bars*: 2 μm. The white arrow, triangle and diamond represented the polar tube, sporoplasm and empty spore coat respectively. The nuclei were stained with DAPI (blue), and the membrane was stained with DiI (red).

By SEM and TEM analysis, the newly forming sporoplasm membrane expanded like a parachute, adhering with the polar tube (Fig. 5A and 5B). Magnification of the sporoplasm showed that it consisted of a thin membrane with a low electron density and the nucleus with high electron density (Fig. 5B). Like blowing soap bubbles, the sporoplasm eventually dropped from the polar tube. There were other compartment-like structures besides the nuclei in sporoplasm, which probably were other intracellular contents transported via the polar tube (Fig. 5C). Moreover, comparatively analysis of absolute quantitative lipidomics between the extruded polar tube and sporoplasm found that the lipid composition of the polar tube and sporoplasm were very similar (Fig. S4), which indicated that the material transported in the polar tube was transferred to the sporoplasm.

**Fig. 5.**
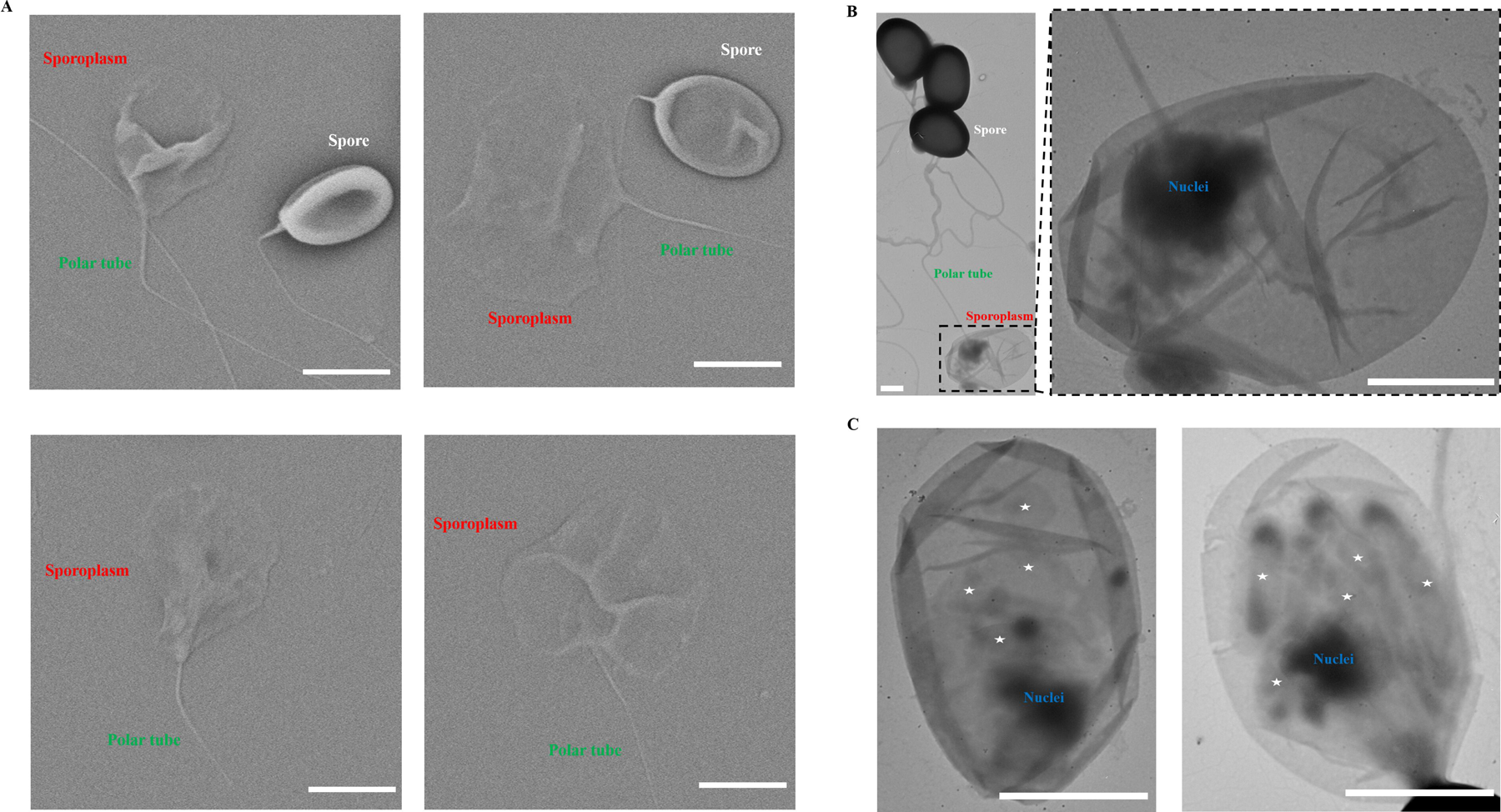
Electron microscope observation the characteristics of *N. bombycis* sporoplasm. (A) SEM analysis of the sporoplasm. *Scale-bars*: 2 μm. (B) and (C) TEM analysis of the sporoplasm. The asterisk represented the membrane structure in the sporoplasm. *Scale-bars*: 1 μm.

### The polaroplast forms the sporoplasm membrane during spore germination

The *N. bombycis* proteins NbTMP1 (Genbank No. EOB13409.1) and NoboABCG1.1 (Genbank No. EOB15202.1) were identified as the surface protein of sporoplasm [39, 40]. In order to further prove the sporoplasm membrane was derived from the polaroplast, we used these two antibodies against sporoplasm surface proteins to label mature spores and found that both NbTMP1 and NoboABCG1.1 localized at the polaroplast in mature spores (Fig. 6A and 6B). After germination, the discontinuous green fluorescence signal of NbTMP1 and NoboABCG1.1 appeared in the extruded polar tube (Fig. 6D and 6E). In *N. bombycis*-infected BmE cells, NbTMP1 and NoboABCG1.1 were also detected on the membrane of sporoplasm (Fig. 6G and 6H). And on the polaroplast, polar tube and sporoplasm membrane, the signal of NbTMP1 and NoboABCG1.1 could co-localize with DiI signal. Based on the above results, we believed that the polaroplast was transported via the polar tube and transformed the sporoplasm membrane.

**Fig. 6.**
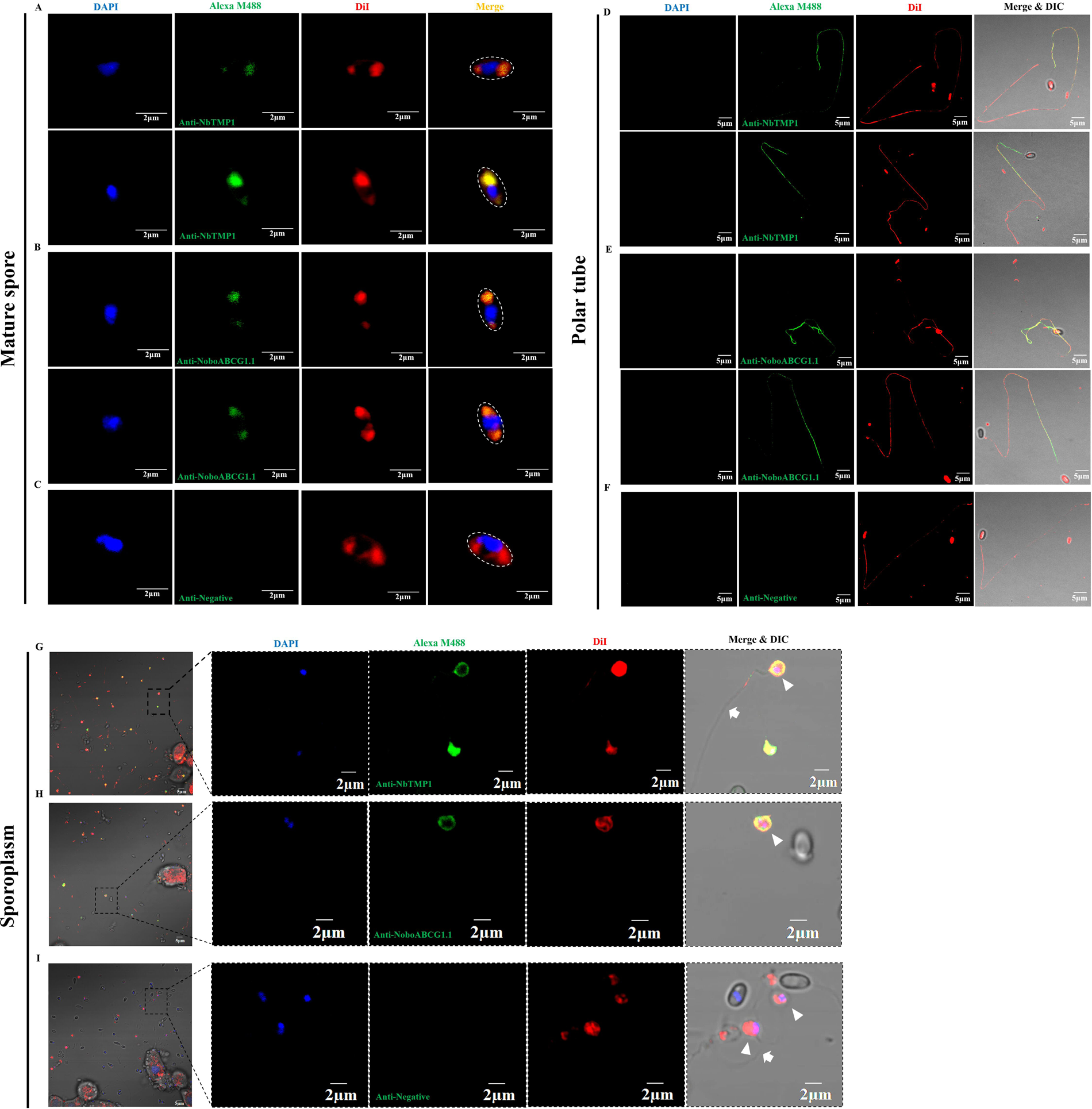
Immunofluorescence assay of NbTMP1 and NoboABCG1.1 localization in the mature spore, extruded polar tube and sporoplasm of *N. bombycis*. (A) and (B) Immunofluorescence assay of NbTMP1 and NoboABCG1.1 localization in the mature spore. (D) and (E) Immunofluorescence assay of NbTMP1 and NoboABCG1.1 localization in the polar tube. (G) and (H) Immunofluorescence assay of NbTMP1 and NoboABCG1.1 localization in the sporoplasm. The mature spore, polar tube and sporoplasm were treated with mouse anti-NbTMP1 and anti-NoboABCG1.1 serum and DiI. (C), (F) and (I) Negative control. The spore, polar tube and sporoplasm were treated with mouse negative serum. the nuclei were stained with DAPI (blue), and the membrane was stained with DiI (red). The white arrow and triangle indicated the polar tube and sporoplasm, respectively. *Scale-bars*: 2 μm and 5 μm.

The origin of the polaroplast, which occupied a large volume in the mature spore, has not been clearly defined. Some researchers have speculated that the polar filament and other infective apparatus components, like polaroplast and PV, are probably the result of Golgi vesicles aggregation [11, 41, 42]. Golgi-tracker green (Beyotime, Shanghai, China) is a Golgi-specific marker, and syntaxin 6 (STX 6) is a protein marker for the Golgi complex [43]. In *N. bombycis*, we found the STX 6 homolog (Genbank No. EOB15057.1) shared 26% homology with human STX 6 (Genbank No. CAG46671.1) and their structure were similar (Fig. S5). Using human STX 6 antibody and Golgi-tracker green labeled mature spores of *N. bombycis*, we found they were localized on the spore polaroplast and partially co-localized with DiI signal of polaroplast (Fig. 7A and 7B). These data are consistent with the hypothesis that the polaroplast derives from Golgi. Additionally, we used Golgi-tracker green and human STX 6 antibody to label the sporoplasm and found that the green fluorescence signal was concentrated on the sporoplasm membrane (Fig. 7C and 7D). These results also support that hypothesis that the polaroplast derived from Golgi finally become the sporoplasm membrane.

**Fig. 7.**
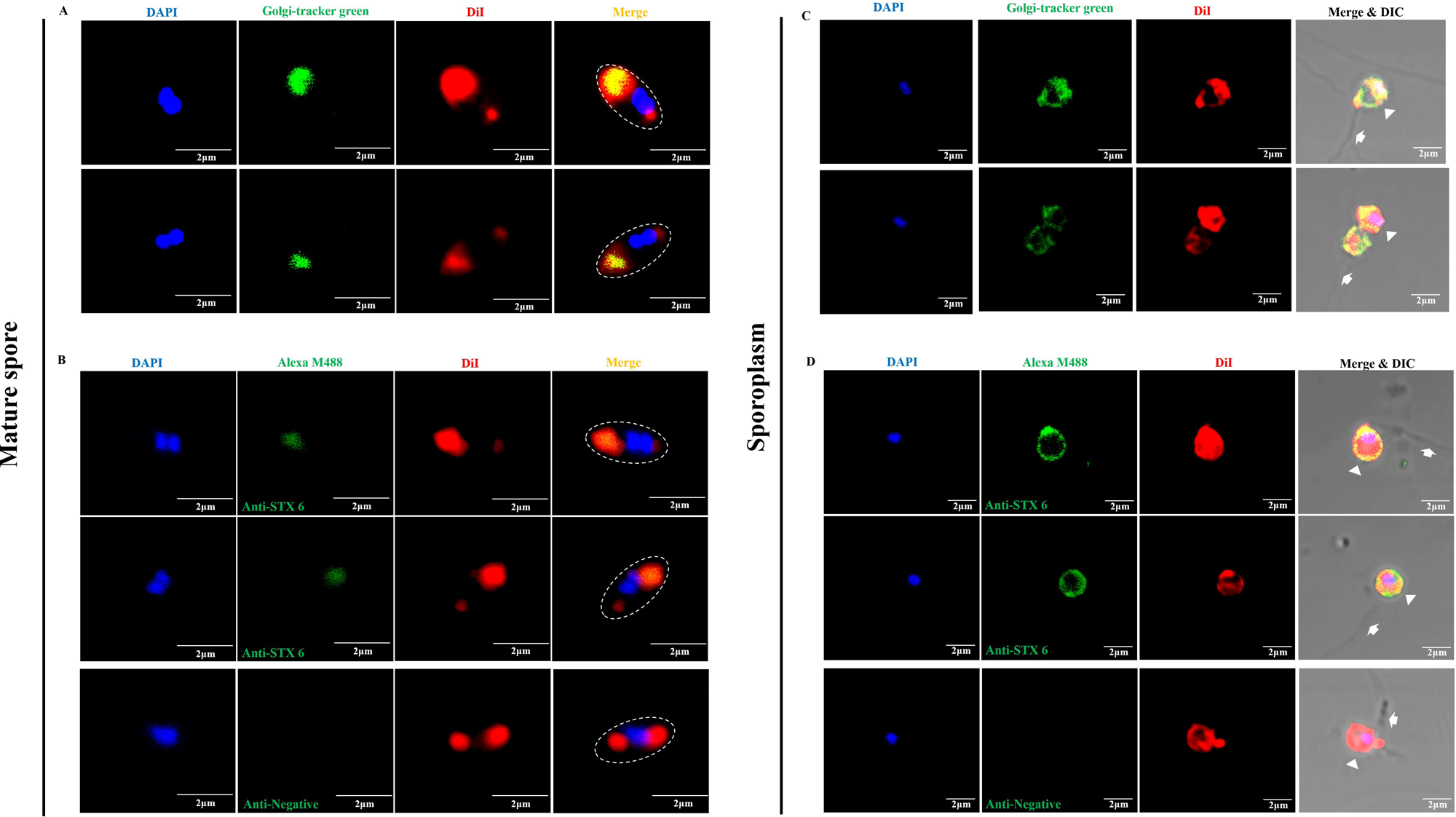
Immunofluorescence assay of Golgi-tracker green and STX 6 in the mature spore and sporoplasm of *N. bombycis*. (A) Fluorescence analysis of Golgi-tracker green in the mature spore of *N. bombycis*. (B) Immunofluorescence analysis of STX 6 in the mature spore of *N. bombycis*. The mature spore was treated with mouse anti-human STX6 serum and mouse negative serum, respectively. (C) Fluorescence analysis of Golgi-tracker green in the sporoplasm of *N. bombycis*. (D) Immunofluorescence analysis of STX 6 in the sporoplasm of *N. bombycis*. The sporoplasm was treated with mouse anti-human STX6 serum and mouse negative serum, respectively. *Scale-bars*: 2 μm.

## Discussion

All members of the microsporidia possess a unique, highly specialized invasion apparatus that includes the polar filament (as polar tube after spore germination), polaroplast and posterior vacuole (PV) [1, 15, 16]. Under appropriate environmental stimulation, the polar tube rapidly discharges out of the spore, pierces the host cell membrane, and acts as a conduit for sporoplasm passage into the new host cell [6, 11]. As a specialized infection apparatus, the structural characteristics of the polar tube are remarkable. The fine structure of polar tube of microsporidium *Anncaliia algerae* was analyzed by Cryo-EM in 2020 [23]. The polar tube surface was pleated and covered with fine fibrillar material. In the polar tube, a variety of transported cargoes were observed, including cylinders, sacs or vesicles filled with particulate material. In this study, we present a germination method for microsporidia spore on Cryo-EM grids to analyze the native structure of extruded polar tube. Distinct with the reported *A. algerae* polar tube, we also found another structure of the polar tube whose surface was no longer fibrillar material, but neatly arranged with bumps of same size like comb teeth, which were speculated PTPs. And this polar tube was filled with uniform electron density materials without any vesicle membrane structures, implying no cargo transportation at this time. Furthermore, comparing the ultrastructure of the polar filament and polar tube by Cryo-EM provides evidence for the hypothesis that the polar filament evaginates during germination.

The polaroplast is a system of membrane-limited cavities in the anterior part of the spore [11, 17, 44, 45]. The expansion of polaroplast results in the disturbed arrangement of the membranous lamellae, thereby increasing turgor within the spore [6, 11, 46–48]. Using two fluorescent probes, N-phenyl-1-naphthylamine (NPN) and chlorotetracycline (CTC), with membrane affinities, fluorescent signals originally in the spore polaroplast were found to transfer to the sporoplasm membrane after spore germination, suggesting that the polaroplast may provide the new plasma membrane for discharged microsporidian sporoplasm [38]. Herein, two antibodies against sporoplasm surface protein of *N. bombycis* was found that they located at the polaroplast in the mature spore, the extruded polar tube and the sporoplasm membrane. In addition, taking advantage of the fact that the polaroplast was derived from Golgi, we used two Golgi markers, Golgi-tracker green and STX 6 protein to label the spore polaroplast and sporoplasm respectively, and found that they were localized in the polaroplast and sporoplasm membrane. Based on the above results, it is further confirmed that the sporoplasm membrane is the polaroplast derived from Golgi. This model overturns the hypothesis that the sporoplasm membrane is the spore plasma membrane [2, 6]. It had been reported that electron microscopy examinations of *Glugea hertwigi* and *Spraguea lophii* spores indicated the presence of the plasma membrane. After the spore germination, this membrane remained in the empty spore coat [38]. In this study, we also demonstrated that the spore plasma membrane and PV remained in the empty spore coat after spore germination. Overall, these data support the hypothesis that the spore polaroplast transforms into the sporoplasm membrane.

The *N. bombycis* sporoplasm is ovate or spherical and about 1.5-2 μm in diameter [35]. Surprisingly, by fluorescence staining, we found that the sporoplasm was not regular spherical at the tip of the polar tube, but gradually changed from an irregular structure to a regular spherical shape over time (Fig. 4). This process was consistent with the model in which the stacked polaroplast vesicles were transported through the polar tube and gradually unfold at the tip of the polar tube, enveloping the nucleus to form the sporoplasm. In the later stages of spore germination, there needs to be a membrane fusion process during the process of sporoplasm shedding from the tip of the polar tube. This process, like blowing soap bubbles, is very similar to the common process of cell refusion after fission [49–51]. A hemi-fused Ω-shaped structure has been observed for the first time in 2016 and is believed to play an important role in cell fusion. This structure was generated by opening and closing of fusion pore on the plasma membrane and the process could be done in a few tens of seconds [52–56]. We also observed a convex punctate structure in the sporoplasm membrane, which might play a key role in membrane closing process (Fig. 4C). Thus, we suspected that a similar process might be occurring when the polaroplast wrapped around the nucleus and fall from the polar tube to form a closed sporoplasm membrane.

The data accumulated to data from the basis for a model for the germination process of microsporidia cargo transport and sporoplasm formation (Fig. 8). When microsporidian mature spore is appropriate stimulated, the posterior vacuole (PV) becomes larger and moves forward, and driven by the PV, the polar tube rapidly evaginates and the front part of the polaroplast is injected into the polar tube first. Then, the spore nucleus is elongated and transported outside through the polar tube. The polaroplast stays at the tip of the polar tube to form a bubble, the nucleus enters the bubble. Finally, the sporoplasm falls off the polar tube. It is noteworthy that both plasma membrane and PV remain in the empty spore coat after spore germination. However, the topology of the polar filament in the mature spore, the relationship between the polar filament and plasma membrane, and the relationship of the polaroplast and plasma membrane, has not yet been determined.

**Fig. 8.**
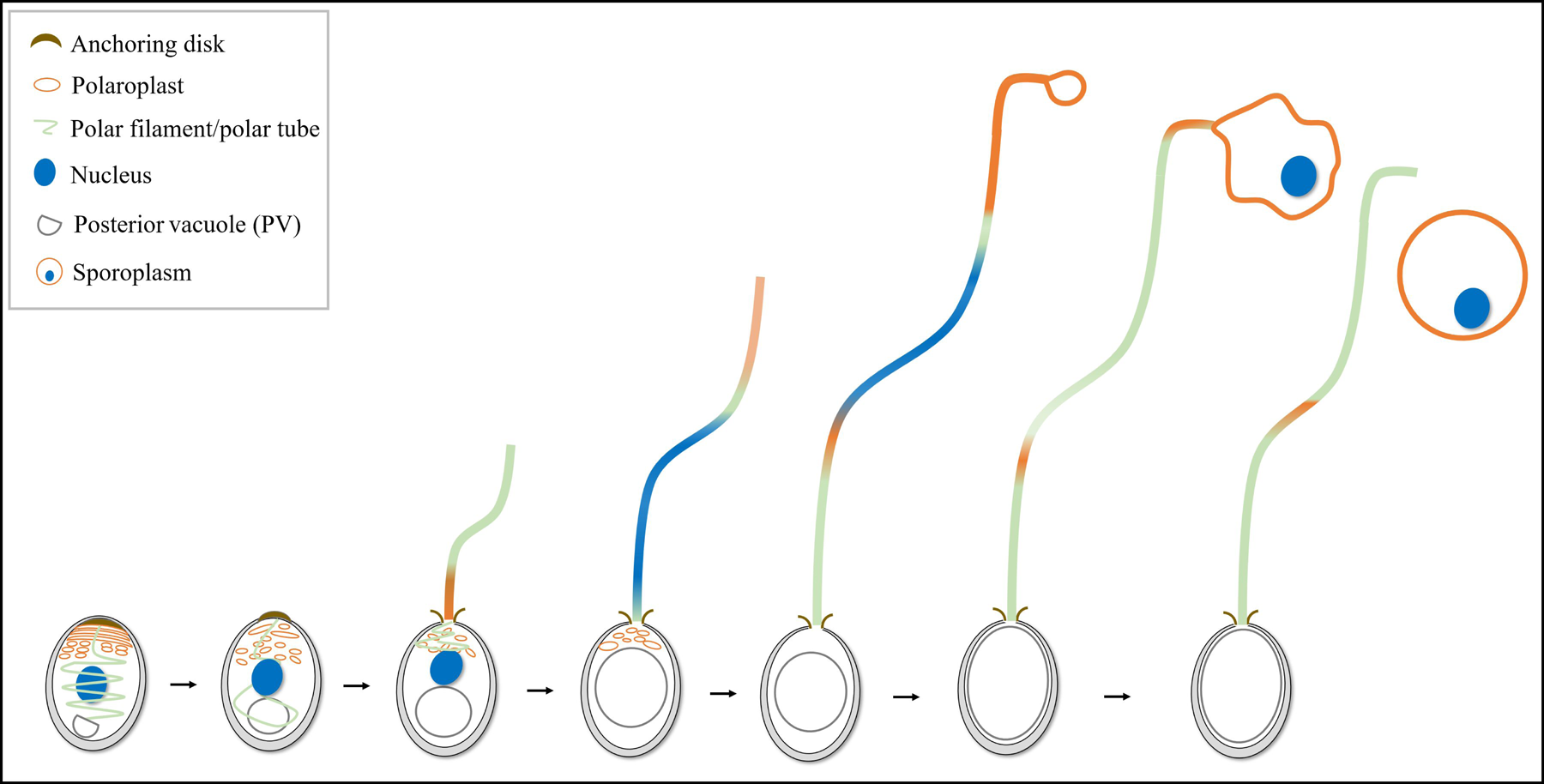
The process of microsporidian germination and sporoplasm formation. When microsporidian mature spore is appropriate stimulated, the posterior vacuole (PV) becomes larger and moves forward. Driven by the PV, the polar filament rapidly evaginates to transform the hollow polar tube, and the front part of polaroplast is firstly injected into the polar tube. Then, the spore nucleus is elongated to transported through the polar tube. The polaroplast stays at the tip of the polar tube to form a bubble, and encloses the nucleus. Finally, the sporoplasm drops off the polar tube.

## Methods

### Preparation and purification of microsporidia spores

Mature spores of *N. bombycis* isolate CQ1, obtained from infected silkworms in Chongqing, China, were obtained from the China Veterinary Culture Collection Center (CVCC no. 102059). Spores were purified using a discontinuous Percoll (Sigma-Aldrich, St. Louis, USA) gradient centrifugation and stored at 4°C until further use.

*E. hellem* spores (*E. hellem* strain ATCC 50504/50451) were collected from infected RK13 (rabbit kidney epithelial cells). Spores were purified from culture media by centrifugation in 70% Percoll at 10,000*g* for 15 min, then washed three times with sterile ddH_2_O, suspended in ddH_2_O, and stored at 4°C until further use [12].

### Cell culture and microsporidian infection

BmE cells (*Bombyx mori* embryo cells) were cultured in Grace medium supplemented with 10% fetal bovine serum (FBS) (Gibco, Thermo Fisher Scientific, Shanghai, China) at 28℃. RK13 cells were purchased from FuDan IBS Cell Center (Shanghai, China) and grown in a complete growth medium supplemented with 10% FBS at 37°C with 5% CO_2_. Vero E6 (African green monkey kidney cells) cell lines were purchased from China Center for Type Culture Collection (Wuhan, China) and were maintained in 10% fetal bovine serum Minimal Essential Medium (MEM) (Thermo Fisher Scientific, Shanghai, China) with penicillin-streptomycin (100 µg/mL, Thermo Fisher Scientific, Shanghai, China) at 37℃ at 5% CO_2_.

*N. bombycis* spores, treated with 0.1 M KOH for 30 s, were added to a 12-well plate containing about 3×10^5^ BmE cells per well to a final ratio of 20:1 (spores: cell) [57]. *E. hellem* spores were a 12-well plate containing added to 5×10^5^ Vero E6 cells according to the 30:1 (spores: cell) ratio [58, 59].

### Immunofluorescence assay (IFA)

The following samples were evaluated as noted below: mature spores, polar tube, polar filament, empty spore coats, sporoplasm, BmE cells infected with *N. bombycis*, and Vero E6 cells infected with *E. hellem*.

*DiI and DAPI staining:* mature spores, polar tube and empty spore coats samples were fixed with 4% *(w/v)* formaldehyde at room temperature for 25 min, and sporoplasms were fixed with 2.5% *(v/v)* glutaraldehyde for 25 min to prevent sporoplasm bust. Then, samples were stained by 1,1’-Dioctadecyl-3,3,3’,3’ tetramethyl indocarbocyanine perchlorate (DiI; Thermo Fisher Scientific, Shanghai, China) for 10 min. Finally, after washing, samples were stained with DAPI (Thermo Fisher Scientific, Shanghai, China) for 15 min.

*DiI, DAPI and Golgi-tracker green staining*: mature spores were fixed with 4% *(w/v)* formaldehyde at room temperature for 25 min, and sporoplasms were fixed with 2.5% *(v/v)* glutaraldehyde for 25 min to prevent sporoplasm bust. Then, samples were stained by Golgi-tracker green (Beyotime, Shanghai, China) for 15 min, and DiI for 10 min. Finally, after washing, samples were stained with DAPI for 15 min.

*Antibody labeling*: 10 μL polar filament samples were added to poly-lysine-coated slides and fixed. Then, the sample was blocked with 10% *(v/v)* non-specific goat serum together with 5% *(w/v)* bovine serum albumin (BSA) in PBST at room temperature for 1 h. Samples were incubated with rab-PcAb-NbPTP1 (1:200 dilution) at room temperature for 1 h. After being washed three times, samples were incubated with Alexa Fluor® 488 conjugate goat anti-rabbit IgG (1:1000 dilution; Thermo Fisher Scientific, Shanghai, China) for 1 h at room temperature. Spore, polar tube and sporoplasm samples were incubated with anti-mouse NbTMP1 (1:200 dilution) and anti-mouse NoboABCG1.1 (1:200 dilution) at room temperature for 1 h, and then samples were incubated with Alexa Fluor® 488 conjugate goat anti-mouse IgG (1:1000 dilution; Thermo Fisher Scientific, Shanghai, China) for 1 h.

Spore and sporoplasm samples were incubated with anti-rabbit STX 6 (1:500 dilution) (ImmunoWay, Jiangsu, China) at room temperature for 1 h, and then samples were incubated with Alexa Fluor® 488 conjugate goat anti-rabbit IgG (1:1000 dilution; Thermo Fisher Scientific, Shanghai, China) for 1 h. After being washed three times, the samples were stained with DiI for 10 min. Following another wash, DAPI was used to stain the samples for an extra 15 min. Finally, all the above samples were examined by the Olympus FV1200 laser confocal microscope (Olympus, Tokyo, Japan).

### Scanning electron microscopy (SEM)

Fifth-instar silkworms were infected with 1×10^4^ spores/mL of virulent *N. bombycis*. The tissues of infected and control silkworms were collected at five days post-infection, respectively, including midgut, fat body, silk gland, testis, ovary and blood. At the late stage of pupae tufting on silkworm, the ovarioles were dissected, and all tissues were washed in ddH_2_O.

Germinated spores were fixed with 2.5% glutaraldehyde (Solarbio, Beijing, China) overnight at 4℃. Then, samples were fixed with 1% osmic acid (Ted pella, Shanghai, China) for 1 h, then were dehydrated using a graded series of ethanol (30%, 40%, 50%, 60%, 70%, 80%, and 90%) for 10 min each and 100% ethanol two times for 15 min each. Following, the samples were dehydrated using tert-butyl alcohol: acetonitrile (2:1 and 1:1), followed by absolute acetonitrile for 10 min each and observed with the scanning electron microscope (Phenom-World BV, Eindhoven, Netherlands).

### Transmission Electron Microscopy (TEM)

*N. bombycis* spores (1×10^5^ spores/mL) were placed on nickel grids (Quantifoil, Beijing, China) for 15 min and germinated in 0.1 M KOH for 30 min at 30°C. Ultrathin sections (70 nm) of mature spore and germinated spore with 0.1 M KOH, and they were prepared as previously described [60] and placed on nickel grids (Quantifoil, Beijing, China). All samples were washed three times by ddH_2_O at room temperature. Then the excess liquid was absorbed and stained in 3% uranyl acetate (Zhongjingkeyi Technology Co., Ltd. Beijing, China), followed by lead citrate (Zhongjingkeyi Technology Co., Ltd. Beijing, China). Images were captured by the Hitachi HT7800 electron microscope (Hitachi, Tokyo, Japan) at 80 kV.

### Cryogenic-electron Microscopy (Cryo-EM)

*N. bombycis* spores (1×10^8^ spores/mL) were placed on grids (Cu R2/1, 200 mesh) (Quantifoil Micro Tools GmbH, Jena, Germany) for 5 min and germinated in 0.1 M KOH for 15 min at 30°C. Cryo-EM grids were prepared by plunge-freezing in liquid ethane on a Vitrobot Mark IV (Thermo Fisher Scientific, USA). The conditions were set as follows: blot time was 2 s; blot force was −2. Then samples were imaged on a 300 kV Titan krios G3i transmission electron microscope (Thermo Fisher Scientific, USA) using a Bioquantum K3 detector (Gatan) with a magnification of 81,000× and the defocus at −3 μm. The EPU software (Thermo Fisher Scientific, USA) was used for automatically sample screening and data collection.

### Quantitative Lipidomics

The polar tube and sporoplasm preparations were pretreated by methyl tert-butyl ether (MTBE) and separated by UHPLC Nexera LC-30A ultra-performance liquid chromatography. Electrospray ionization (ESI) positive and negative ion modes were used for detection, respectively. The samples were separated by UHPLC and analyzed by mass spectrometry with Q Exactive series mass spectrometer (Thermo Fisher Scientific, Shanghai, China). The mass charge ratios of lipid molecules and lipid fragments were collected by the following method: 10 fragment maps were collected after each full scan (MS2 scan, HCD). The MS1 has a resolution of 70,000 at M/Z 200, and the MS2 has a resolution of 17,500 at M/Z 200. LipidSearch was used to carry out peak recognition, peak extraction and lipid identification (secondary identification) of lipid molecules and internal standard lipid molecules. Precursor tolerance: 5 PPM, product tolerance: 5 PPM, product ion threshold: 5%. This project was completed in Shanghai Applied Protein Technology Co. Ltd. (Shanghai, China).

## Data Availability

All other relevant data are within the manuscript and its Supporting information files.

## Acknowledgements

We sincerely thank Ms. Shiyi Zheng and Mr. Hongyun Huang for kindly providing anti-NbTMP1 serum. We appreciate the efforts of Mr. Junrui Guo and Ms. Xue Wang in their assistance with cell culture. Thanks to Dr. Qiang He for his friendly help in the sporoplasm purification and observation. We thank Ms. Chunxia Wang and Yan Zhou for assistance with the TEM sample preparation and observation, and Ms. Lei Liu for assistance with Cryo-EM sample preparation.

## Author Contributions

Q.L. conceived, designed and performed experiments, analyzed data and wrote the manuscript. Y.Q.C., L.Q., L.Y.H., Z.K.X., H.J.L., X.L.H. and C.X.L. performed experiments, and analyzed data. X.Z.M. and J.C. provided advices and experimental materials. B.H., Q.T.S., and L.M.W. gave advice and edited the manuscript. Z.Y.Z. supervised research. M.X.L. and G.Q.P. supervised research, designed experiments and edited the manuscript.

## Supplementary Information

**Fig. S1 TEM analysis of the extruded polar tube of *N. bombycis*.**

(A) Spore was germinated by KOH, and the polar tube was released. *Scale-bars*: 5 µm. The white arrow in the enlarged image points to the discontinuous high electron density material of the polar tube. *Scale-bars*: 1 µm. (B) The curved hooks at the anterior end of polar tube. *Scale-bars*: 500 nm.

**Fig. S2 Structure characteristics of the polar filament in *N. bombycis*.**

(A) Immunofluorescence analysis the structure characteristics of polar filament. The polar filament was treated with rabbit anti-NbPTP1 serum (green) and DiI (red). The nucleus was labeled with DAPI (blue). The white arrow represented the polar filament and the white triangle represented the DiI-labeled membrane structure. *Scale-bars*: 5 μm. (B) Cryo-EM observation the structure characteristics of polar filament. *Scale-bars*: 100 nm.

**Fig. S3 TEM analysis of the mature spore and germinated spore of *N. bombycis*.**

(A) Mature spore. *Scale-bars*: 1 μm. (B-E) Spores germination by 0.1 M KOH. The white dotted line part is the posterior vesicle. *Scale-bars*: 1 μm. (F) The polaroplast in the mature spores. *Scale-bars*: 500 nm. The white arrow represented the polar filament, the white dotted line represented the posterior vesicle, and the yellow dotted line represented the polaroplast.

**Fig. S4 Absolute quantitative lipidomics analysis of the polar tube and sporoplasm in N. bombycis.**

**Fig. S5 Characterization of the STX 6 in *N. bombycis*.**

(A) Sequence alignment analysis of human STX 6 (Genbank No. CAG46671.1) and *N. bombycis* STX-like protein (Genbank No. EOB15057.1). The red and green boxes represented the functional domains and transmembrane domains of human STX 6 and STX-like protein respectively. (B) Three-dimensional structure prediction of human STX 6 and STX-like protein.

